# Creation and manipulation of bipartite expression transgenes in *C. elegans* using phiC31 recombinase

**DOI:** 10.1101/2024.03.01.583017

**Authors:** Michael L. Nonet

## Abstract

Bipartite expression systems are widely used in model organisms to express specific gene products in a cell-specific context. They are typically encoded as two independent, unlinked transgenic insertions: a driver and a reporter. Herein, I outline an efficient method named <u>recombination-mediated h</u>omolog <u>e</u>xchange (RMHE) that uses genetically encoded phiC31 recombinase to convert driver and reporter transgenes integrated at the same genetic position from a *trans* configuration where one component is on each chromosome, to a *cis* configuration where the driver and reporter are on the same chromosome. This approach is based upon the development of a set of 3’ *attB* tagged driver lines and 5’ *attP* tagged reporter lines. This genetic based approach leverages both the power of combinatorial re-assortment of drivers and reporters and the simplicity of single locus genetics. I also describe a novel microinjection-based approach named <u>recombination-mediated i</u>ntegration (RMI) that utilizes the individual *attB* driver and *attP* reporter lines as landing site for the phiC31 recombinase mediated integration of whole driver and reporter plasmids into the existing reporter and driver lines, respectively. Thus, this work outlines both a novel genetic based and a novel microinjection-based method to create cis-linked driver/reporter pairs. These new tools increase the utility of bipartite systems for *C. elegans* genetics by reducing the complexity of reporter system segregation in crosses and thus can greatly simplify the use of bipartite reporter systems during genetic analysis.

## Introduction

Bipartite expression systems are powerful transgenic tools that can be used to express genetically encoded sub-cellular markers, to manipulate cellular function using RNAi and similar approaches, and to facilitate cell-type specific biochemistry using bioID style approaches (Dietzl et al., 2007; Jin et al., 2012; Wang et al., 2017; Branon et al., 2018; Kamiyama et al., 2021). They are based on two distinct transcription units, one expressing a transcription factor called a ‘driver’ and one expressing a reporter responsive to the driver (Brand and Perrimon, 1993). The benefits of bipartite systems include the ability to amplify the signal of weak promoters and the flexibility provided by separating the components. In most systems, these two transcription units are independently derived transgenic insertions, though it is possible to create insertions that contain both a driver and a reporter (Distel et al., 2009). Distinct reporter and driver insertions can be combined to greatly expand the utility of the individual transgenes. In *Drosophila melanogaster*, drivers and reporters are typically kept as independent stocks and crossed to analyze a double heterozygote carrying both transgenes (Caygill and Brand, 2016). However, in *C. elegans* this is not practical for several reasons. First, *C. elegans* is a self-fertile hermaphrodite and strains are usually passaged by passive self-crossing of hermaphrodites. Second, balancers chromosomes are much less well developed in *C. elegans* than in *Drosophila*. Third, few well tolerated visible markers are available to track chromosomes and combinations of these visible markers are often not feasible to score or maintain together. Fourth, performing forward genetic screens and RNAi screens is often not practical using out crosses due to the rapid life cycle of worms. Thus, in practice, working with bipartite systems in *C. elegans* is often best performed in animals homozygous for both the reporter and the driver unless lethality or sterility are caused by toxic or developmental consequences of the specific bipartite pair (Wei et al., 2012; Wang et al., 2017; Nonet, 2020). Construction of such dual homozygotes is relatively straightforward and easily accomplished in two generations (6 days). Introduction of mutations into these backgrounds is also feasible with outcrosses followed by backcrosses. More difficult is utilizing additional bipartite tools in such backgrounds. For example, split-driver systems (Wang et al., 2018; Luan et al., 2020), dual driver/repressor systems which require 3 transgenes (Suster et al., 2004; Wei et al., 2012), and using two distinct bipartite reporter systems in concert, which requires 4 transgenes. While crossing two strains which contain distinct drivers and reporters is straightforward, deriving a strain homozygous for three or four markers is more complex. Performing these manipulations while introducing a lesion of interest is even more complicated. Due to these significant constraints, I sought a practical means to simplify manipulation of bipartite systems in worms.

Site specific recombinases offer the potential to simplify the creation of bipartite driver-reporter strains by catalyzing recombination between transgenic insertions positioned at the same site in the genome. In such a scheme, a driver construct carrying a recombination site on the 3’ of the insertion and a reporter carrying a compatible recombination site on the 5’ of the insertion are recombined from *trans* to *cis* by the recombinase, creating a novel chromosome containing both the reporter and the driver (**Fig. 0**). I refer to this general approach as <u>Recombination-Mediated H</u>omolog <u>E</u>xchange (RMHE). In principle, both reciprocal recombinases like Cre and FLP and non-reciprocal recombinases like phiC31 and Bxb1 could be used to create these novel transgenes.

**Figure 0.**
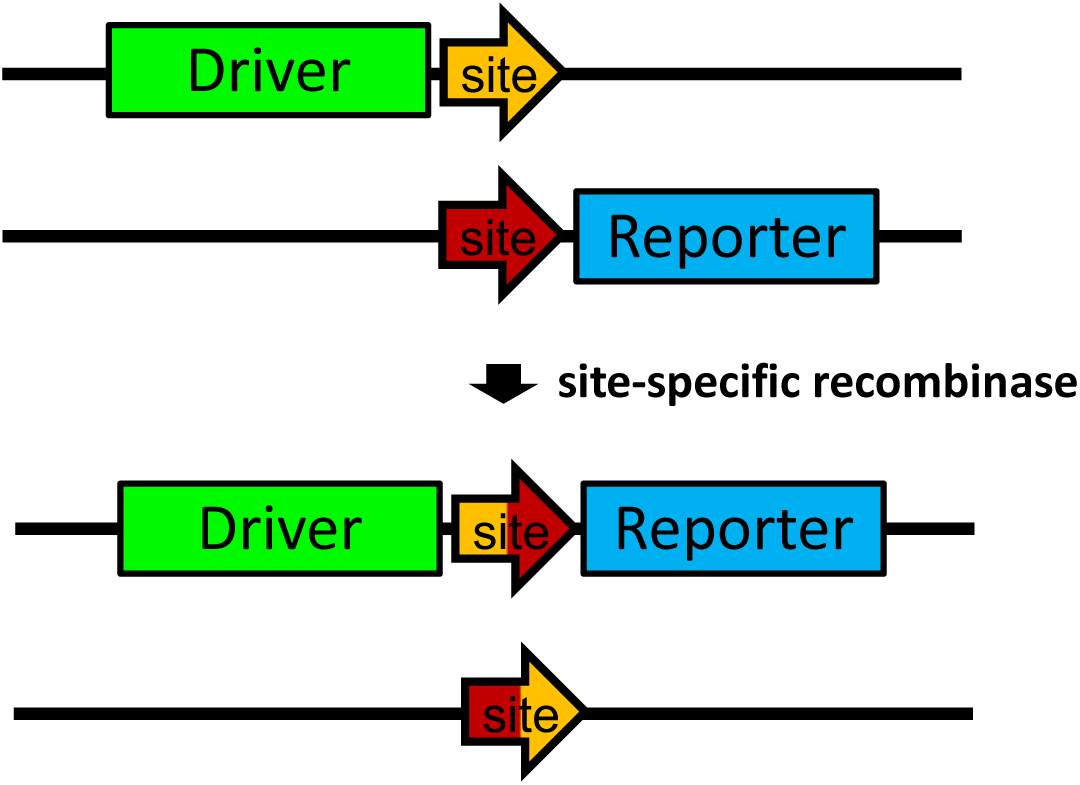
Recombination of bipartite transgenes from *trans* to *cis* configuration. Schematic of recombinase-based reassortment of two transgenic insertions integrated at the same position on a chromosome from *trans* to *cis* using a site-specific recombinase. The two insertions are differentially tagged on the 5’ and 3’ of the insertion with recombinase sites and are recombined via site-specific recombination in the germline to create the novel *cis* configuration along with another chromosome containing only the recombination site. The approach is shown for a non-reciprocal recombinase such as phiC31 but is conceptually similar to that for a reciprocal recombinase such as FLP.

Here I demonstrate that in *C. elegans*, phiC31, but not FLP, recombinase will catalyze recombination events between homologous chromosomes and can efficiently mediate the conversion of *trans* driver/reporter heterozygotes into *cis* driver-reporter combinations on the same chromosome. In addition, I develop and demonstrate the utility of RMHE integration vectors compatible with rapid recombinant-mediated cassette exchange (rRMCE) integration methods to simplify creation of driver and reporter transgenes. I show these transgenes can easily be recombined to create stable homozygous driver::reporter lines. I demonstrate the utility of this system for creating traditional organelle markers, TIR1 expressing lines for analysis using conditional auxin inducible degradation methods (Zhang et al., 2015), and for creating cell specific GFP1-10 expressing lines for visualizing protein localization at endogenous levels using a NATF approach (He et al., 2019). Furthermore, I demonstrate that phiC31 recombinase can also be used to integrate *att* site containing plasmids into *attB* driver or *attP* reporter strains at high efficiency by germline injection into strains harboring an *att* site and expressing both *phiC31* and FLP. These tools provide practical means of creating bipartite reporter constructs that are linked but retain the combinatorial advantages of having independent driver and reporter transgenes.

## Results

### Tests for FLP-mediated recombination

I first designed transgenic animals to assess the efficiency of FLP recombinase to perform RMHE in *C. elegans* using transgenic animals created using the recombination-mediated cassette exchange (RMCE) integration method. Since RMCE utilizes FLP and *FRT* sites to create the initial transgenes, the advantage of using *FRT* sites as the sites for RMHE is that one does not need to introduce an additional recombinase to catalyze the reaction. A set of distinct *FRT* sites that do not recombine with each other have been previously characterized (Schlake and Bode, 1994; Turan et al., 2010). Among these I chose to introduce *FRT13* sites into the 3’ end of driver constructs and the 5’ end of reporter constructs. This created a pair of transgenes that could be recombined at the *FRT13* site to switch the two transgenes from the *trans* to *cis* configuration (**Fig. 1A**). *Trans* heterozygotes containing both a tet^OFF^ driver *FRT13* and a *FRT13 tetO* reporter insertions in a *bqSi711* germline FLP expressing background were crossed with a *myo-2p*::GFP marked male, then subsequent cross progeny were characterized (**Fig. 1B**). In two cases using different pairs of transgenes, no recombination events were observed among 1381 cross progeny examined. This suggests that recombination between *FRT13* sites at similar positions in homologous chromosomes does not occur at an appreciable frequency (**Fig. 1C**). To assess whether recombination was occurring at a low frequency or in somatic tissue, I performed PCR analysis of DNA isolated from the progeny of individual *trans*-heterozygotes. PCR amplification did not detect the expected recombinant *FRT13* chromosome despite easily detecting the expected product from non-transgenic wild type animals and both non-recombinant chromosomes products (**Fig. 1D**). The genetic and molecular data suggest that recombination between *FRT13* sites on distinct chromosomes is rare.

**Figure 1.**
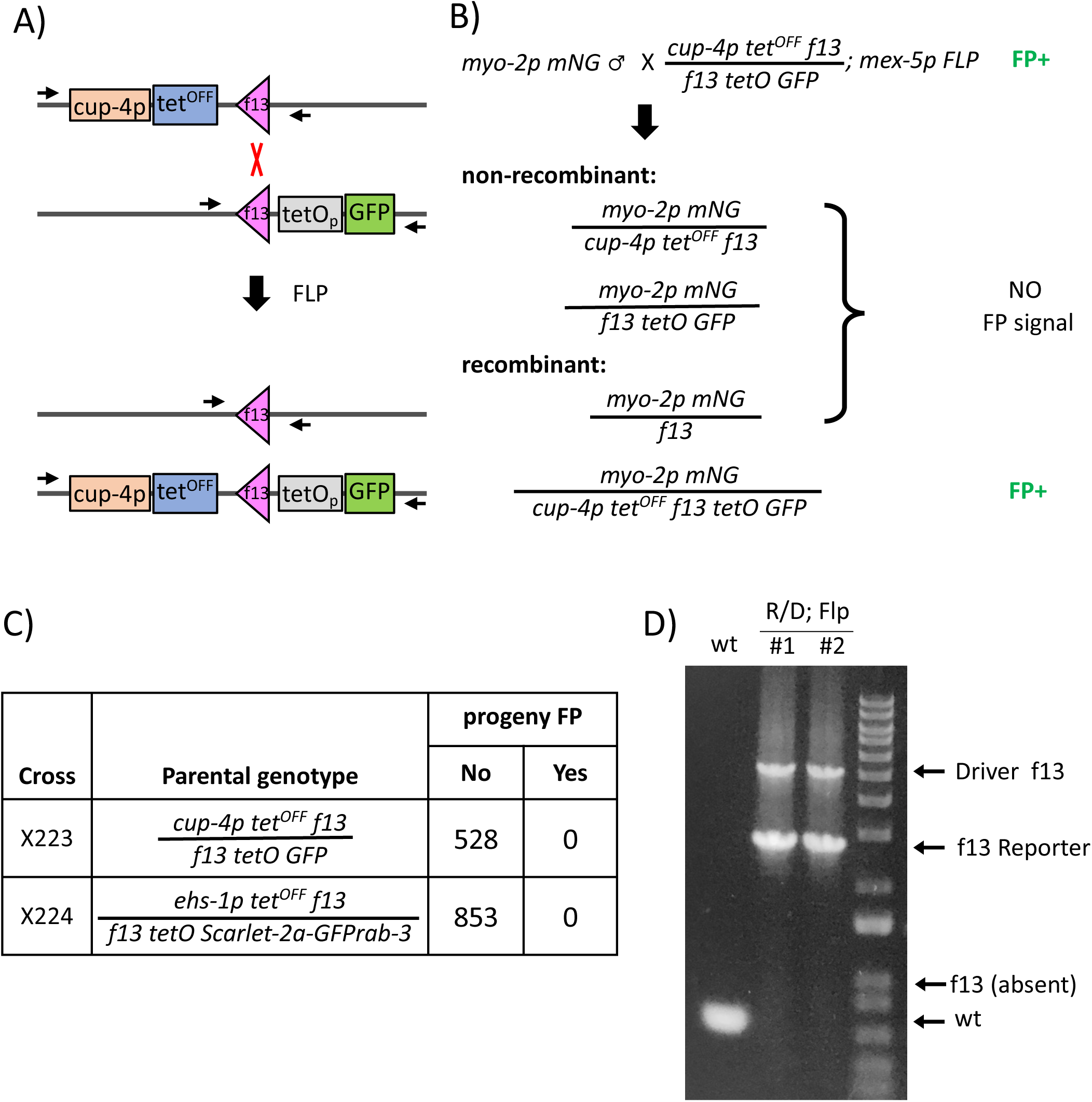
Test for FLP-mediated homolog exchange. **A**) Schematic of the methodology used to test the ability of FLP to perform RMHE. Shown are the structure of the chromosomes of a *trans* heterozygote containing *FRT13* (f13 labeled pink triangle), tagged *cup-4p tet*^OFF^, and *tetOp GFP* transgenic insertions along with the expected structure of the chromosomes after recombination by FLP. **B**) Schematic of the genetic cross used to screen for the presence of recombined chromosomes. Note that the heterozygous *mex-5p FLP* chromosome present in the cross progeny is omitted from the diagram for clarity. **C)** Summary table of the strains used to quantify the frequency of FLP-mediated recombination. **D)** PCR analysis of pooled progeny of two individual *cup-4p tet*^OFF^ / *tetOp GFP*; *mex-5p FLP a*nimals and the wild type as a control. The positions of a PCR control wild type non-transgenic chormosome, of the parental driver and reporter chromosomes and the expected position of the f13 recombined chromosome lacking the driver and reporter are marked with arrows. The position of the two oligonucleotides used for PCR is depicted in panel A. Strains used: NM1 wt, NM5382 *jsSi1602/ jsSi1598 II; bqSi711 IV,* NM5407 *jsSi1615/ jsSi1618* II; *bqSi711* IV. See Table S1 for a list of the strains used and their complete genotypes.

### Testing if pairing effects recombination

FLP has successfully been used to recombine homologous chromosomes in somatic tissue in Drosophila and this event is the basis for the MARCM (Mosaic Analysis with a Repressible Cell Marker) technique (Potter et al., 2010). However, in addition to MARCM occurring in the soma, the two chromosomes in such experiments are also perfectly paired in the recombination interval with the unique genetic elements of each chromosome being distant from the *FRT* sites. By contrast, in the experiments described above, the two chromosomes are not likely to be well paired at the *FRT* site due to lack of homology between the two transgenes around the FRT sites. Thus, I designed two chromosomes which minimizes differences between the two chromosomes and in which the two chromosomes could pair perfectly within 1220 bp 5’ and 444 bp 3’ of the *FRT13* sites (**Fig. S1A**). In addition, the *FRT13* sites used are complete sites (48bp) that contain three 13 bp repeats, matching those used in initial studies characterizing FLP RMCE (Schlake and Bode, 1994). This is in contrast to minimal sites containing only two repeats that have become the standard in the field because the third repeat is dispensable for insertion and excision (Ringrose et al., 1999). One insertion consisted of a *mec-4* synthetic promoter (*mec-4Sp*) driving tet^OFF^, a *FRT13* site and a *tetO* promoter (*tetOp*) driving GFP-his-58. The other consisted of the *mec-4Sp* driving tet^ON^, a *FRT13* site and a *tetOp* driving an *mCherry-his-58* fusion (**Fig. S1**). Even in this case, recombination between the chromosomes was not detected in 252 F1 cross progeny (**Fig. S1B**). PCR experiments were not performed since the long homologous region adjacent to the *FRT13* site would very likely lead to cross annealing of PCR amplicons from the two chromosomes, yielding to hybrid PCR products even without recombination occurring. I conclude that FLP is unable to efficiently catalyze recombination between homologous chromosomes in the germline of *C. elegans*.

### Expression of phiC31

phiC31 is another recombinase that has been widely used to manipulate genomes. To test if phiC31 could catalyze chromosome exchange I first used RMCE to create a transgene in which expresses a *C. elegans* codon optimized phiC31 with a C-terminal nuclear localization signal and three synthetic introns using a design guided by approach used in Drosophila (Bischof et al., 2007). Although the use of phiC31 for RMCE has been described in *C. elegans* (Yang et al., 2022), my studies and constructions were initiated independently and prior to public disclosure of that method (Nonet, 2021). *phiC31* was expressed under the control of the *mex-5* promoter and the *glh-2* 3’ UTR and contained a downstream sl2 mNG cassette to facilitate genetic tracking of the transgene. The phiC31 transgene, *jsSi1623*, expresses mNG at comparable levels to strains expressing FLP in a similarly designed transgene at the analogous position in the *C. elegans* genome on Chr IV (Macías-León and Askjaer, 2018). Another transgene, *jsSi1822,* expressing the same construct was also integrated on Chr II.

### Tests for phiC31-mediated recombination

To test for the ability of phiC31 to recombine *attB* and *attP* sites in *C. elegans* I developed *attB* and *attP* containing transgenes that could be distinguished in the same animal. I created *attB mec-4p nls-mNG* transgenes and *mec-4p Scarlet-his-58 attP* transgenic animals using RMCE (**Fig. 2A**). Crosses were performed to create *attB/attP* trans-heterozygotes in a background homozygous for a germline phiC31 expressing transgene. Progeny were screened visually for recombination events and recombinants were identified. Surprisingly, the *cis* recombinants expressed the GFP marker in the pharynx (**Fig. 2B)**, while the *trans* heterozygote animals did not which made the recombinants easy to identify in heterozygotes using pharyngeal expression as a dominant marker. This ectopic expression is reminiscent of the crosstalk observed when RMCE transgenes are produced that contain multiple closely spaced transcription units with distinct expression profiles (Nonet, 2023). Using this assay, I quantified the recombination frequency and determined that 20.5% (46/224) of chromosomes derived from an *attB/attP trans* heterozygote converted from the *trans* to *cis* configuration (**Fig. 2C**). To confirm that recombination was occurring, I performed PCR analysis to detect the small *attL* reciprocal product that occurs during phiC31 mediated recombination. This product was present in DNA isolated from the progeny of trans heterozygotes carrying phiC31 but absent from animals lacking the recombinase **(*Fig. 2D*).**

**Figure 2.**
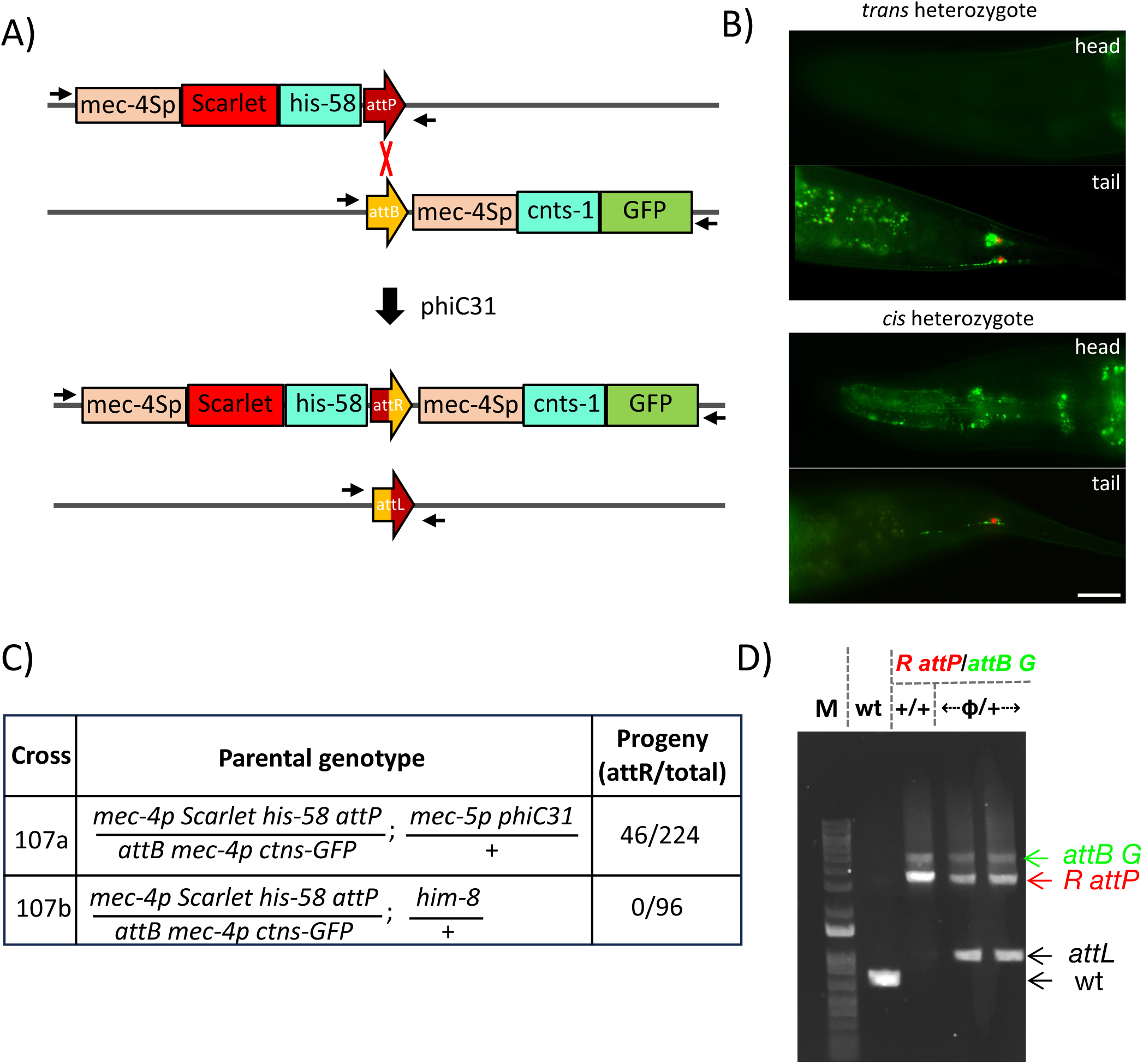
Test for phiC31-mediated homolog exchange. **A)** Schematic of the methodology used to test the ability of phiC31 to perform RMHE. Shown are the structures of the chromosomes of a *trans* heterozygote containing an *attP* tagged *mec-4p Scarlet-his-58* transgenic insertion and an *attB* tagged *mec-4p cnts-1-GFP* insertion along with the expected structure of the chromosomes after recombination by phiC31. **B**) Images of the head and tail regions of a *trans-*heterozygote parent animal carrying both unrecombined chromosomes and a cis-heterozygote carrying one recombinant chromosome and one *mec-4p Scarlet-his-58* parental chromosome. **C**) Summary table of the self-crosses used to quantify the frequency of phiC31-mediated recombination. **D**) PCR analysis of pooled progeny of individual *mec-4p Scarlet-his-58 attP / attB mec-4p ctns-1-GFP*; *mex-5p phiC31/+* animals, and *mec-4p Scarlet-his-58 attP / attB mec-4p ctns-1-GFP* animals lacking phiC31 as well as the wild type as controls. The position of the two oligonucleotides used for PCR is depicted in panel A. See Table S1 for a list of the strains used and their complete genotypes. Scale Bar 20 µm.

To further characterize the phiC31 recombination frequencies I created tet^OFF^ driver constructs containing a 3’ *attB* site and *tetO* reporter constructs carrying a 5’ *attP* and integrated these using RMCE on chromosome II. After introducing the phiC31 expressing transgene into the *attB tetO* reporter strain, crosses were performed to assess the frequency of recombination both in hermaphrodites and males carrying a single copy of the phiC31 expressing transgene (**Fig. S2A**). When driver *attB*/reporter *attP trans* heterozygous males were crossed to *unc-119* animals, *cis* recombined chromosomes were observed in 3.8% (38/1000) of cross progeny while when *myo-2p mNG* males were crossed to *trans*-heterozygous hermaphrodites, *cis* recombined chromosomes were observed in 9.2% (92/1000) cross progeny (**Fig. S2A**).

To confirm that the ability of phiC31 to recombine homologous chromosomes was generally applicable across the genome rather than restricted to specific hot spots, *attB* and *attP* transgenes were inserted on Chr I and the recombination frequency observed in these crosses was quantified. On Chr I, recombinants were obtained in 14.5% (188/1301) of cross progeny of hermaphrodite and 7.4% (101/1368) of male trans-heterozygotes carrying a single copy of phiC31 (**Fig. S2B,C**). Recombinants were also readily obtained from *attP* and *attB* containing insertions on Chr IV and V though the frequency was not quantified.

### Minimal *attB* and *attP* sites are functional

The initial phiC31 recombination experiments were performed using 285 bp *attB* and 221 bp *attP* regions that extend beyond the minimal required sequences for efficient intramolecular recombination in *E. coli* (Groth et al., 2000). Transgenes containing only those minimal sequences were constructed and tested for recombination. Recombination was observed in 5.7% (23/400) of cross progeny from a hermaphrodite, and 3.5% (7/200) of cross progeny from a male *trans*-heterozygote carrying one copy of phiC31 when both minimal *attB* and *attP* sites were used (**Fig. S3A**) and 5.4-6.2% (31/500 and 27/500 for hermaphrodites) and 1.9%-2.5% (7/374 and 2/79 for males) were obtained when a minimal *attP* was recombined with a larger *attB* (**Fig. S3B**).

In summary, recombination between *att* sites positioned at analogous positions on homologous chromosomes can be catalyzed by phiC31. The frequency of this event is dependent on both the sex of the germline and the size of the *att* sites and is likely also influenced by the copy number of the phiC31 expressing transgene.

### (r)RMCE compatible RMHE integration tools

To facilitate the development of transgenic lines for RMHE, I developed Golden Gate (GG) compatible integration vectors for both the RMCE and rRMCE integration methods (Nonet, 2020; Nonet, 2023). The first vectors developed to create RMHE compatible driver and reporter lines are based on a RMCE vector (**Fig. S4A**) that relies on a *sqt-1* Self Excision Cassette (SEC) to identify insertion events (Nonet, 2020). These tools yield unmarked driver and reporter lines (**Fig. S5A**). To make RMHE compatible transgenes using rRMCE, which also permits *cis* marking of insertions, I modified pHygG and pHygR series vectors (Nonet, 2023) by incorporating *attB* and *attP* sites. These vectors (**Fig. S4B, Table S1** and **Supplemental methods** for details) enable the creation of driver *attB* and reporter *attP* lines that are also marked with nls-GFP or nls-Scarlet under the control of a cell specific promoter such as *myo-2p* (pharyngeal muscle), *mec-4p* (TRNs) or *cup-4p* (coelomocytes) to track the driver and reporter in crosses (**Fig. S5B**). After performing RMHE to create cis-recombined driver-reporter combinations, the insertion remains marked with the cell specific FP associated with the driver and, if desired, this marker and the *Hyg^R^* cassette can subsequently still be excised using Cre.

### tet^OFF^ driver lines

To demonstrate the utility of RMHE, my lab generated numerous tet^OFF^ constructs with cell-specific promoters in RMHE vectors (see *Table S1* for a complete list). The targeting plasmids were integrated into the genome and characterized for expression in both a *trans* and *cis* genomic conformation by recombining the driver *attB* transgenes with an *attP tetO nls-GFP* reporter transgene (**Fig. 3A**). Most of the promoter tet^OFF^ vectors directed strong nls-GFP expression in a cell specific pattern similar to that previously reported for the promoter (**Fig. 3B-G**). However, a few expressed in broader patterns than the published description of cell-types obtained using direct promoter-FP fusions (*mir-228p* in **Fig. 3H** and *dpy*-*7p* **Fig. S6A**). For example, mir-228p expressed much more broadly than the glial-specific signal previously reported (Fung et al., 2020). I suspect that weak expression of tet^OFF^ in certain cell types is amplified into detectable signal from the *tetO 7X nls-GFP* reporter. Most promoter::tet^OFF^ drivers behaved similarly in the *trans* and *cis* configuration when expressing the reporter specifically in the expected cell type. However, some exhibited ectopic expression in the *cis* configuration that was not observed in the *trans* configuration (**Fig. 3I**). For example, pgp-12p expressed stochastically in intestinal cells in addition to expressing in the excretory cell (Zhao et al., 2005). I performed experiments to address this issue.

**Figure 3.**
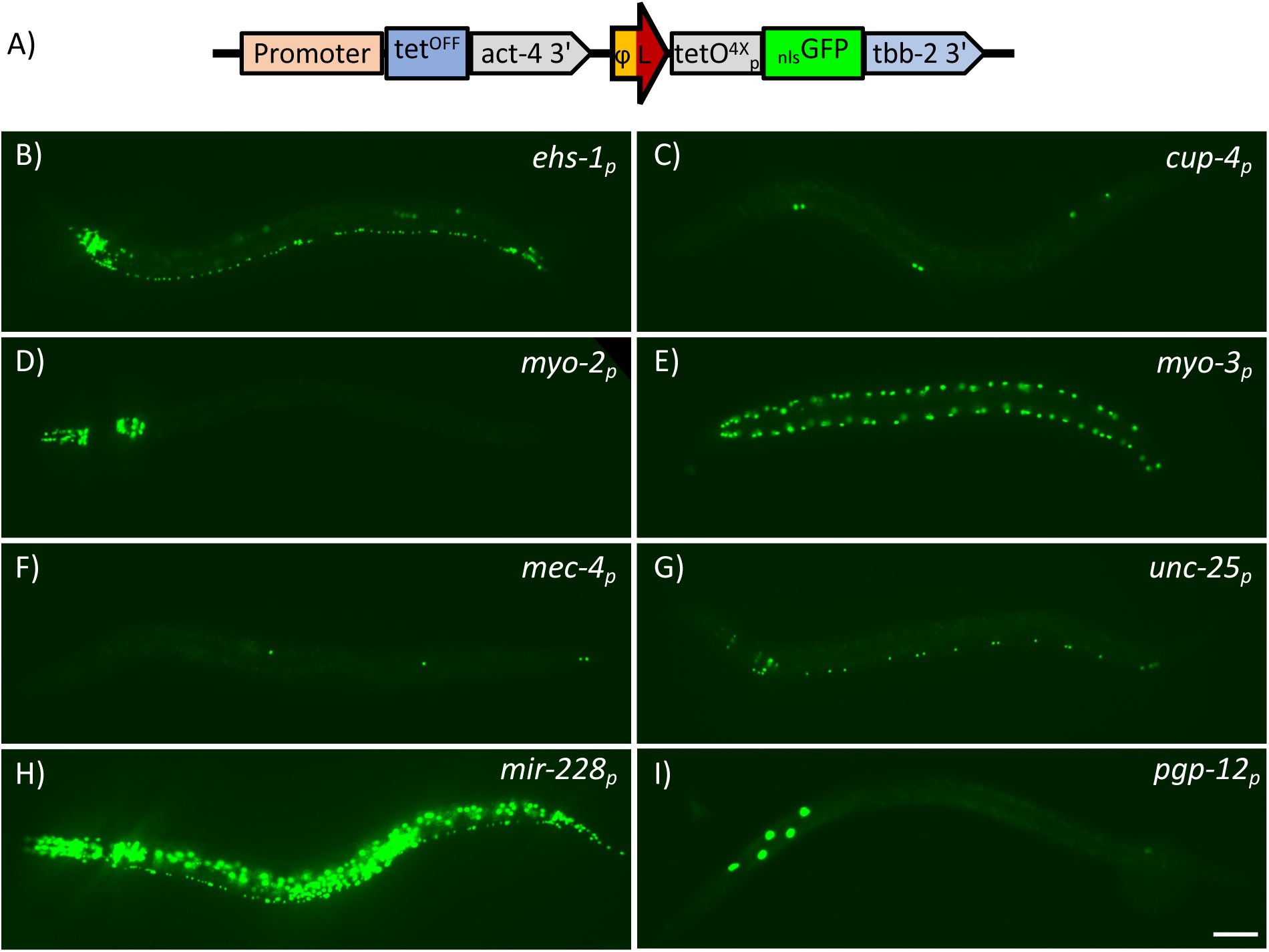
Expression pattern of c*is* linked *tet^OFF^* driver *tetO_p_* reporter transgenes. **A)** Diagram of the structure RMHE-derived *cis* linked *tetOFF* driver *tetOp* reporter transgenes. ***B-I***) Images of L4 animals carrying *tetOFF* driver *tetOp* reporter transgenes driven by distinct promoters listed in the top right of each image. See Table S1 for a list of strains used and complete genotypes. Scale bar 50 µm.

### *Cis* vs. *trans* analysis

In the initial experiments to test for phiC31-mediated recombination, I observed unexpected differences between the expression pattern of two transgenes when placed in *cis* versus in *trans* (**Fig. 2B**). Similar phenomena were also observed after recombining tet^OFF^ driver and tetO_p_ nls-GFP reporter lines. Specifically, in *trans* the expression pattern of the GFP reporter was that expected of the cell specific promoter used to express the driver (**Fig. 4A**). However, for certain driver-reporter combinations, when recombined in *cis,* expression was observed in the pharyngeal nuclei in the posterior of the pharynx (**Fig. 4B**). To identify the cause of this ectopic expression, several additional lines were created. Both *nhx-2p* and *mec4p* lines exhibited the unusual expression after recombination from *trans* to *cis* suggesting it was independent of the promoter being used (***Fig. 4B* and *Fig. S7A*).** The ectopic expression of the tet^OFF^ lines was abolished by the addition of doxycycline indicating that the influence was through ectopic expression of the driver, rather than driver-independent effects of an enhancer acting on the reporter basal promoter (**Fig. 4G**). In addition, an *nhx-2p* GAL4 attL *UAS 11X nls-GFP* line also exhibited the same ectopic expression phenomenon suggesting it was not a feature of a particular bipartite system (**Fig. 4C**). Insertions on Chr I and Chr II both exhibited ectopic expression in the pharyngeal gland cell nuclei, but insertions of Chr I also showed expression in the rectal gland cell and the pharyngeal/intestinal valve cell (**Fig. S7A**). The differential patterns observed at the two insertion sites suggests that at least some of the influences are derived from the genomic environment of the landing site. One obvious difference between *cis* and *trans* configurations is the structure of the *att* site between the two transgenes, which is changed after recombination. To eliminate the possibility that *attL* was acting as a cryptic promoter or enhancer, constructs containing both a driver and a reporter separated by either a periodic A_n_/T_n_-clusters region (PATC)-rich sequence (**Fig. 4D**), or an inter promoter region (**Fig. S7B**) were constructed and integrated into the same genomic position as the RMHE constructs. The two insertions that removed the *attL* site still showed similar ectopic expression as the RMHE insertions indicating that *attL* is unlikely to be responsible for the ectopic expression. Increasing the spacing between the reporter and driver reduced but did not eliminate the signal (**Fig. S7D**). Finally, several commonly used 3’ UTRs, including the *tbb-2* 3’ UTR, have been shown to have promoter enhancing activity (Knoebel et al., 2024). Thus, I replaced the 3’ UTRs in the driver and reporter and created additional RMHE lines. Replacing the *tbb-2* 3’ UTR of the reporter with a *let-858* 3’ UTR eliminated the ectopic expression (**Fig. 4E**) and replacing the *act-4* 3’ UTR of the driver with a *ter222* synthetic 3’ UTR, a synthetic terminator that includes the tbb-2 3’ UTR (Knoebel et al., 2024), partially restored the ectopic expression (**Fig. 4F**). In summary, the source of this ectopic expression appears to be the commonly used *tbb-2* 3’ UTR, though the experiments performed do not provide satisfactory rationale for why the ectopic expression is *cis*-transgene specific.

**Figure 4.**
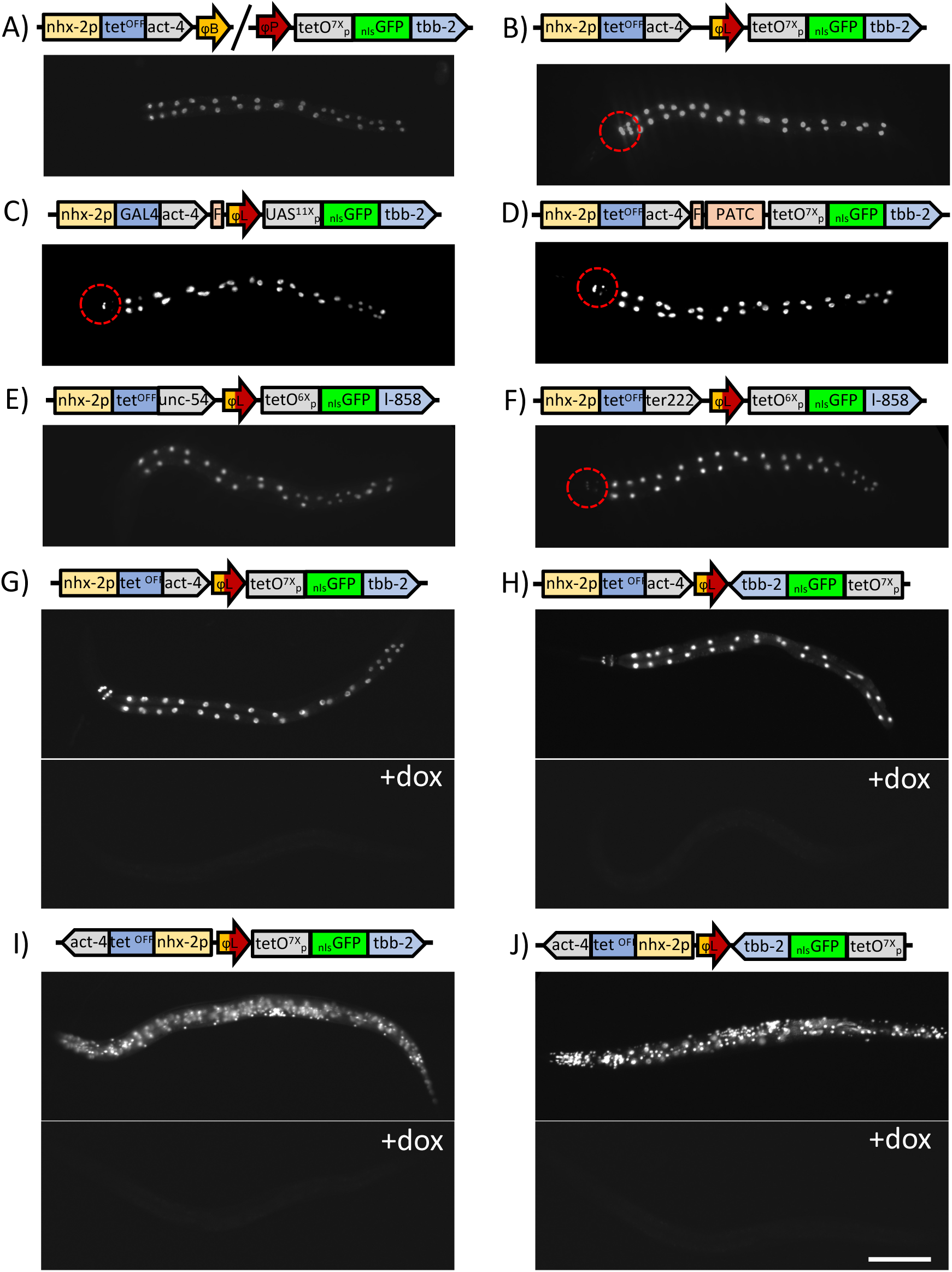
*cis* vs. *trans* and orientation dependent influences on bipartite reporter expressio. **A-B**) Images of L4 animals showing a comparison of the expression pattern of an *nhx-2p tetOFF* driver and *tetOp nls-GFP* reporter pair in **A**) *trans* and in **B**) *cis.* The red circles denote the ectopic pharyngeal expression. **C-H**) L4 images of *cis nhx-2p tetOFF* driver and *tetOp nls-GFP* reporter pair with substitutions to identify the source of the pharyngeal background. **C**) Replacing *tet^OFF^* and *tetO* with *GAL4* and *UAS* reduced but did not eliminate the background. **D)** Replacing the *attL* site with a periodic A_n_/T_n_-clusters region (PATC) did not eliminate the background. **E**) Replacing the *act-4* 3’ UTR and *tbb-2* 3’ UTR with the *unc-54* and *let-858* 3’ UTRs, respectively, eliminated the background. **F**) Replacing the *act-4* 3’ UTR with *ter222*, a synthetic terminator that contains the *tbb-2* 3’ UTR, the *rpl-2* 3’ UTR, and the *rps-2* 3 ‘ UTR in tandem partially restored the pharyngeal background. **G-J**) Tests for the effects of orientation of the *tet^OFF^* driver and the *tetOp* reporter. With the transcription orientation of the driver facing toward the reporter (**G-H**), the orientation of the reporter had little effect on specificity of expression and a modest effect on background in the pharynx. By contrast, when the transcription orientation of the driver was facing away from the reporter (**I-J**), expression of the reporter was virtually non-specific with expression occurring at high levels in many different tissues regardless of the orientation of the driver. Both the intestinal signal and the non-specific signals were conditional with expression being completely eliminated by the addition of doxycycline (**G-J**, bottom panels). See Table S1 for a list of strains used and complete genotypes. Scale Bar for all images 100 µm.

### Orientation tests

In building *cis* linked driver and reporter transgenes, there are four ways the driver and reporter could be oriented with respect to each other. I constructed *nhx-2p* and *mec-4p* tet^OFF^ drivers in both orientation and recombined them with *tetO* GFP reporters in both orientations. Reversing the orientation of the reporter such that it transcribed towards the driver has minimal effect on the expression pattern of the cis-driver-reporter transgene (**Fig. 4G, H and S7C, D, G,H**). However, unexpectedly, positioning the driver in an orientation divergent from the *tetO* reporter, regardless of the orientation of the reporter, yielded high non-specific expression in many tissues making transgenic drivers in that orientation unusable as cell specific transgenic tools (**Fig. 4I, J and S7I, J**). Increasing the spacing between the driver and reporter in these orientations did not significantly reduce the level of ectopic expression (**Fig. S7E, F**). All of the ectopic expression was completely eliminated by adding doxycycline (**Fig. 4I, J and S7I, J).** I hypothesize this orientation creates a feedback loop in which low level expression of the tet^OFF^ transgene is directed by the *tetO* reporter. Whether this occurs by failure of RNA polymerase II termination or engagement of the basal promoter elements of the driver promoter by the *tetO 7X* enhancer (or a combination of the two) remains unclear. The lack of orientation specificity of this effect is likely the result of bi-directional transcription that occurs at *tetO* promoters (Knoebel et al., 2024).

### *tetO* reporter lines

In addition to creating cell-specific driver lines, my lab also created a large set of reporters lines (see **Table S1** for a complete list) including organelle specific markers, tools for monitoring and disrupting synaptic activity (Davis et al., 2008; Nguyen et al., 2017), and other *C. elegans* genetically encoded tools such as TIR1 and GFP1-10 transgenes for AID and NATF analysis (Zhang et al., 2015; He et al., 2019). Most reporters were expressed under the control of a *tetO 4X mec-7bp* promoter. This promoter initially was my view of the optimal signal to noise ratio for expressing in TRNs using a *mec-4p* driver. However, subsequent more in-depth characterization of parameters influencing expression levels in bipartite reporter systems indicates this is likely a sub-optimal, yet reasonably functional, promoter for many other cell types (Knoebel et al., 2024). I concentrated on building sub-cellular reporters that replicated previously vetted tools (Rolls et al., 2002; Yu et al., 2008; Thomas et al., 2019). To validate the reporters, I recombined them with a driver to visualize the expression pattern as a simple means of confirming similar behavior to the published reporters. Different strategies were used to identify animals carrying the recombined chromosomes depending on the structure of the reporters and drivers (**Fig. S8**). The expression level of the reporters varied substantially with some being relatively weak and others being exceedingly bright (**Fig. 5 and S6**). In general, transgenes which contained codon optimized genes or encoded a reporter in which the first coding portion of the reporter (i.e., GFP) was codon optimized, expressed very well. In contrast, those with sub-optimal (often native) codon usage expressed less well. The vast majority localized as expected. However, in some cases overexpression obviated the intended utility of the tool such as with the microtubule (MT) +TIP *ebp-2-mNG,* typically used to visualize the plus ends of microtubules, which expressed so strongly that the entire MT cytoskeleton was well labelled (**Fig. S6D**). Thus, some tools will need to be optimized based upon the cell specific expression levels obtained by altering the number of *tetO* sites, the basal promoter or the 3’ UTR (Knoebel et al., 2024).

**Figure 5.**
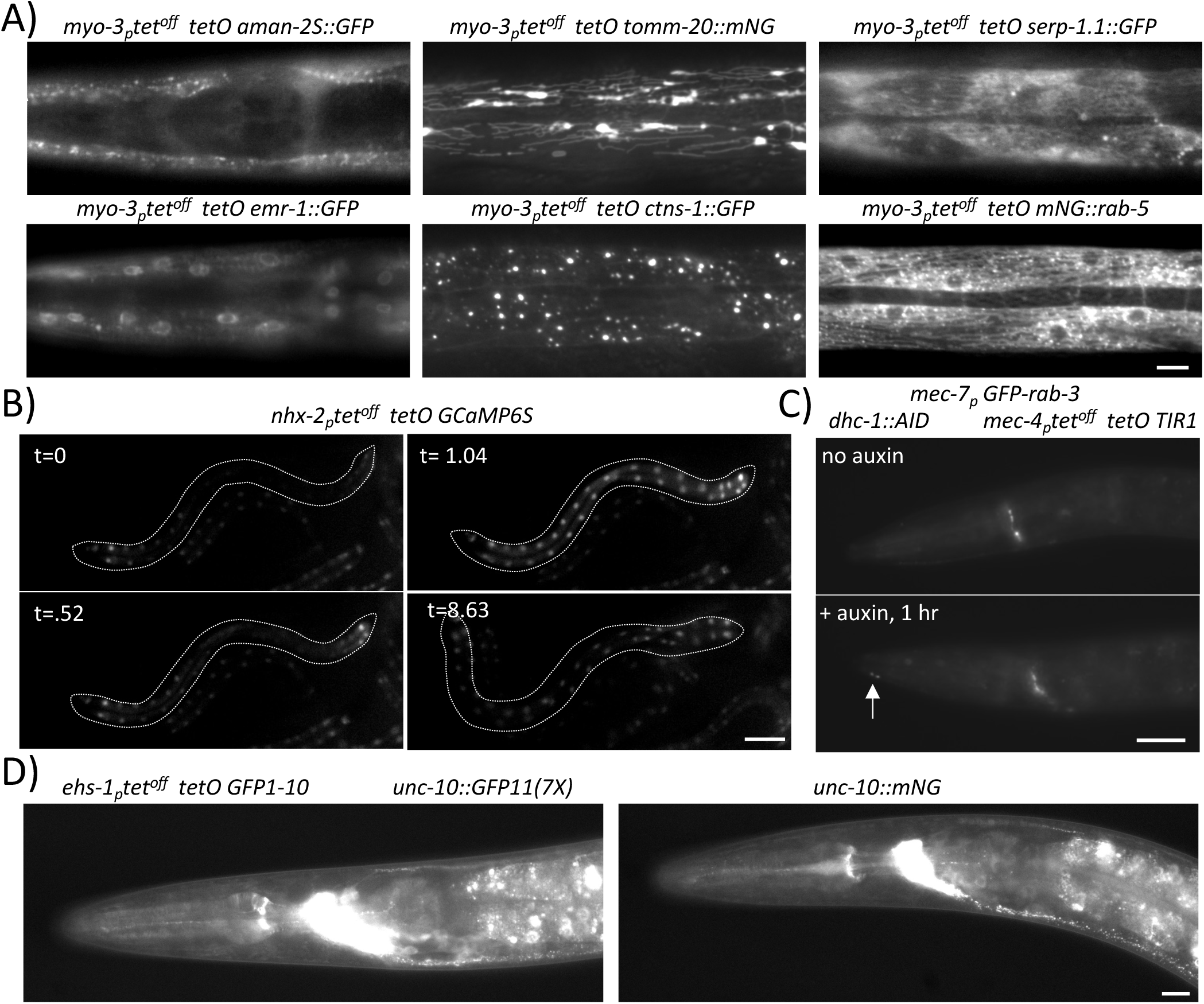
Validation of bipartite reporter system derived organelle markers, sensors, and tools. **A)** Images of L4 muscle showing the localization pattern of various organelle markers as labeled: (left top) head muscle with Golgi labeling, (left bottom) head muscle with nuclear envelope labeling, (middle top) body wall muscle with mitochondrial labeling, (middle bottom) body muscle with lysosomal labeling, (right top) body wall muscle with rough ER labeling, (right bottom) body wall muscle with early endosome labeling. Scale Bar 10 um. **B)** Images from a movie of L3 animals expressing the calcium sensor GCaMP6S to visualize the intestinal calcium wave during defecation (Teramoto and Iwasaki, 2006). The anterior of the animal shown is on the left of the image. Time in seconds is listed in the top left of each panel. The entire movie is provided as a supplement. Scale Bar 50 µm. **C)** Images of the head and midbody of L3 *dhc-1::AID* animals expressing *TIR1* and *GFP-rab-3* in TRNs before (top) and after 1 hr. (bottom) in the presence of 0.5 mM auxin. Arrows point to the end of the ALML axon, which is barely detectable before addition of auxin, but accumulates GFP labeled synaptic vesicle precursors when the dynein motor is degraded. This phenotype parallels that observed in *dhc-1* mutants (Koushika et al., 2004). Scale Bar 20 µm. **D)** Images of the head of L4 *unc-10::GFP11(7X)* animals expressing pan neuronal *GFP1-10* to visualize synaptic active zones (left) and an L4 *unc-10::mNG* animal for comparison. See Table S1 for a list of strains used and complete genotypes. Scale Bar 10 µm.

### Construction of a dual FLP phiC31 transgene

To further demonstrate the utility of *attB-attP* recombination and to simplify the RMHE pipeline after construction of *attB* transgenes using RMCE, a strain expressing both FLP and phiC31 was constructed using RMHE. Such a dual recombinase expressing strain is desirable to permit RMCE derived *attP* insertions to be directly crossed to *attB* promoter strains that already express phiC31. *bqSi711* expresses FLP under the control of *mex-5p* on Chr IV. To construct the desired transgenes, I integrated an *attB* site just 3’ of the *bqSi711* insertion using CRISPR (**Fig. S9A**) and then created a 5’ *attP mex-5p phiC31* transgene using RMCE (**Fig. S9B**). The two strains were then crossed, and individual progeny were cloned. Twenty animals were screened by PCR to identify animals containing a RMHE recombined chromosome (**Fig. S9C**) resulting in the isolation of three recombinants, one of which was used to isolate a homozygous *jsSi1784* transgene expressing FLP, phiC31 and mNG in the germline under control of *mex-5p*. In addition to creating a desirable recombinase tool, this experiment demonstrates that existing transgenes can be tagged with an *att* site and recombined with other transgenes integrated at the same position in the genome.

### Selecting for RMHE using drug resistance

RMHE is efficient enough that even if a visual screen is not feasible for identifying recombinants, PCR can be used reasonably effectively to identify the desired recombinant. Nevertheless, in certain situations recombination frequencies might be unexpected low or the resulting recombinants might be unhealthy and difficult to recover. Therefore, vectors that permit the selection for recombination events were also created. rRMCE vectors that contain a split Neo cassette with adjacent *att* sites were constructed (**Fig. S10A**). The orientation of the split Neo cassette was set such that it transcribed toward the *Hyg^R^* selection cassette to minimize the potential for crosstalk. In addition, a *lox2722* site was placed in each vector so that the Neo selection cassette could be excised after performing RMHE, if desired. To test the system, *mec-4p* tet^OFF^ was introduced into the *attP* vector and *tetO mNG* into the *attB* vector. The two plasmids were integrated on Chr II using rRMCE, and then recombined via a cross through *a* phiC31 expressing line (**Fig. S10B**). Neo^R^ animals expressing mNG were isolated and homozygosed. These animals express mNG in TRNs as expected, but the expression level was lower than typically observed with the *mec-4 tet^OFF^* transgene. The resulting transgene was crossed through a Cre expressing line. Animals derived from the Cre cross homozygous for a recombined chromosome with both the *cis* marker and the *Neo^R^* cassette excised now express comparably to on *mec-4p* tet^OFF^ transgene (**Fig. S10B**). These Neo selection vectors offer a methodology to create transgenic constructs using RMHE that might be very sick or lethal by using the conditional nature of the tet system.

### Direct <u>R</u>ecombination-<u>M</u>ediated Integration (RMI) into existing *att* marked drivers and reporters

In some cases, one has developed a reporter or a driver line, but one does not have the appropriate partner for RMHE. The efficiency of RMHE suggested that it might be possible to integrate directly into a genomic phiC31 *att* site using a targeting plasmid rather than a performing RMHE with a previously integrated line. Indeed, Yang et al. (2022) have previously used phiC31 to perform RMCE. I developed an additional integration method, which I have called <u>recombination-mediated i</u>ntegration (RMI), where an entire targeting plasmid is integrated by a single recombination event. To test this approach, I developed RMI integration vectors that contained an *att* site, a *FRT* site, a MCS compatible with *SapI* Golden Gate cloning, and a *lox2722* flanked *Hyg^R^*, *Neo^R^* or split NeoC selection cassette for selection for inserts (**Fig. S11**). Two initial experiments using a split Neo vector were performed because in the split Neo system the isolation of Neo^R^ animals is strongly indicative of a recombination event at the landing site. The results suggested that RMI was very efficient (see **supplemental methods** for details). However, the split Neo selection is not compatible with the myriad of reporter and driver lines developed for RMHE herein. Thus, additional tests were performed using existing reporter and driver lines.

To assess the frequency of RMI, a set of distinct promoter::tet^OFF^ combinations were inserted into an *attB* RMI vector, and various *tetO 7X nls-FP* and *tetO 7X FP* combinations were inserted into the *attP* RMI vector. After introducing the *jsSi1784* phiC31 and FLP expressing transgene, groups of plasmids were co-injected in the *attB* and *attP* lines (**Fig. 6, S12, Table S2**). The co-injected plasmids were chosen specifically to be distinguishable as insertions without requiring molecular analysis. Three days after the injection, hygromycin or G418 was added to select for insertions. As was the case with the initial tests, I observed much higher rates of resistant animals from plates with 2 or 3 injected animals than typically obtained with RMCE. The vast majority of plates contained >100 viable healthy animals by three days after drug addition and the plates starved by day 5 or 6; two days earlier than one would typically begin to observe individual L4 animals with homozygous insertions using RMCE. In the initial experiment. I co-inject 2 targeting plasmids, and both were integrated in all 4 plates of two injected P0s (**Table S2**). In additional experiments I co-injected 6 distinct plasmids. In these injections, I recovered multiple (3-6) different types of insertions from injections of 2 (occasionally 3) P0s (**Table S2**, **Fig. S12C**). The conversion of a *cis* marker from Scarlet to mNG (**Fig. S12A, B**), the similarity of expression levels and patterns in independent isolates, and PCR analysis all indicate that the insertions are occurring at the intended *att* site (**Fig. S12D**). Notably, different plates often showed a predominance of one or two insertions and a lower frequency of other insertions. Overall, the rate of insertion of RMI appeared much higher than RMCE, and I conservatively estimate it is at least an order of magnitude higher. Supporting that conclusion, I identified numerous F2 animals that were heterozygous for two different insertions indicating that distinct insertions had occurred in the sperm and oocyte of individual animals. As noted above, Individual plates were dominated by different insertion lines suggesting that early jackpot insertions occur. These early events make quantification of the exact frequency difficult from these experiments.

**Figure 6.**
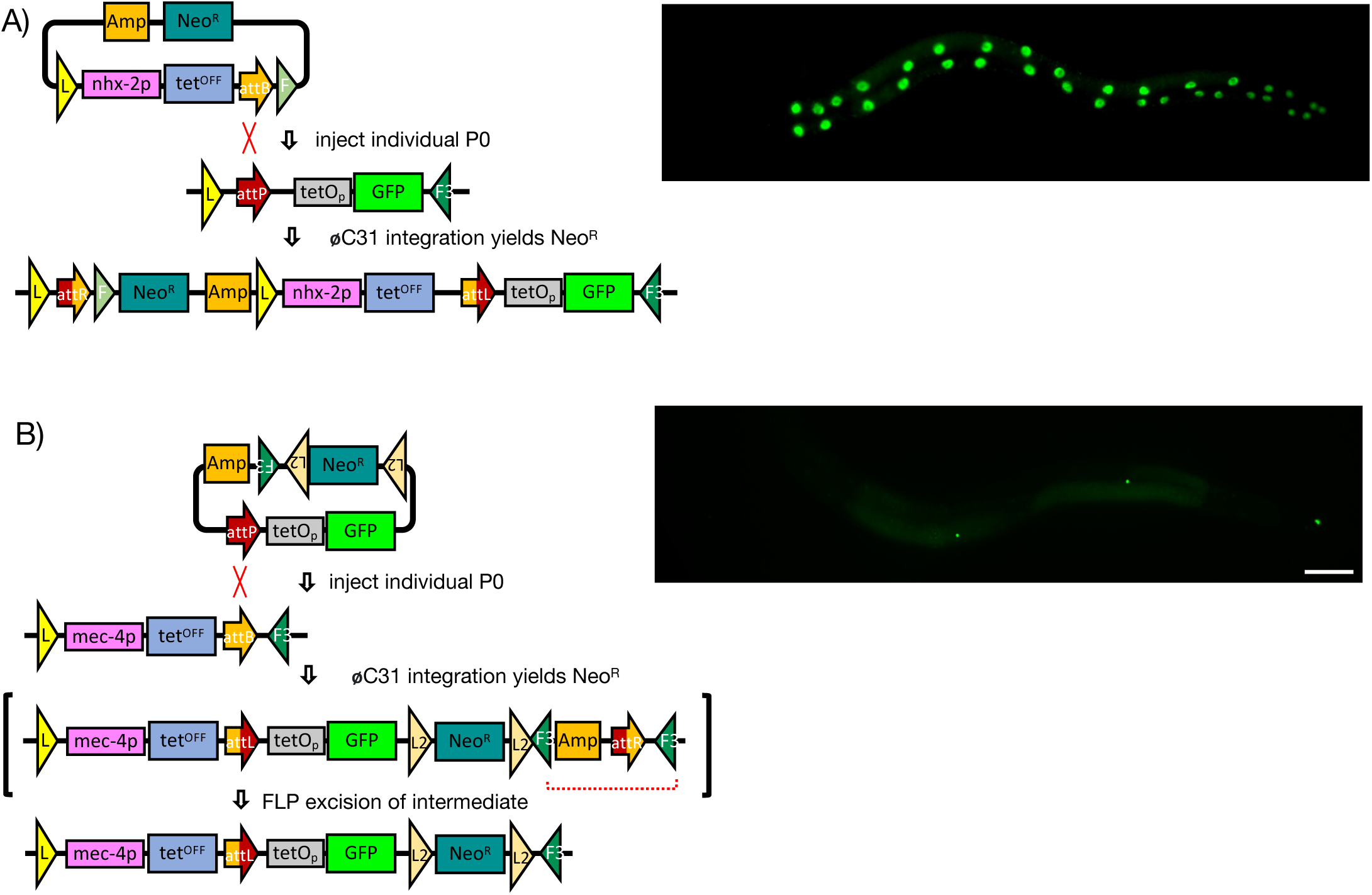
Overview of recombination-mediated integration (RMI) of entire plasmids. **A-B**) Schematics of integration plasmids and *att* landing sites used to assess the efficiency of RMI. The unlinked *jsSI1784* phiC31 and FLP expressing transgene is omitted for clarity. On the right are images of L4 animals homozygous for the resulting transgenes. The light diffuse germline mNG signal derives from the *jsSi1784* transgene still present in the strains. *attB* and *attP* sites are shown as orange and red arrows and the recombined *attL* and *attR* sites as dual color arrows based on the recombination event. Yellow triangles labeled ‘L’ represent *loxP* sites and light and dark green triangles labeled ‘F’ and ‘F3’ represent *FRT* and *FRT3* sites, respectively. Integration of the *attP* plasmid yields an unstable product containing the vector backbone and an *attR* site flanked by a pair of *FRT3* site which is efficiently excised by FLP. The orientation of the triangle represents the orientation of these recombination sites in the construct. See Table S1 for a list of strains used and complete genotypes. Scale Bars 50 µm.

To quantify the insertion frequency, young adult worms were injected with a library of plasmid pAttBFGNeoN14 (complexity ∼ 10,000). After selection for Neo^R^ for 2 generations, the F3 generation of animals was collected. Genomic DNA was prepared from the mixed sample, then insertion events were amplified by PCR and the resulting fragments were purified and subject to nanopore sequence analysis (**Fig. 7A, B**). It was assumed that individual reads of the mixed PCR product are proportional to the frequency of distinct insertions each containing a unique bar code. Analysis of the reads revealed that multiple different integrations occurred in 9 of 10 animals. In all of these cases, a few distinct insertions were most abundant, but in addition, from 2 to 15 additional insertions were obtained at lower frequency (**Fig. 7C**). These data indicate that phiC31 RMI insertion frequency is at least an order of magnitude higher than rRMCE where the average insertion rate is approximately 1 per 2 injected animals (Nonet, 2023). The increase in frequency raises the possibility that insertions might occur at other locations in the genome besides the intended site.

**Figure 7.**
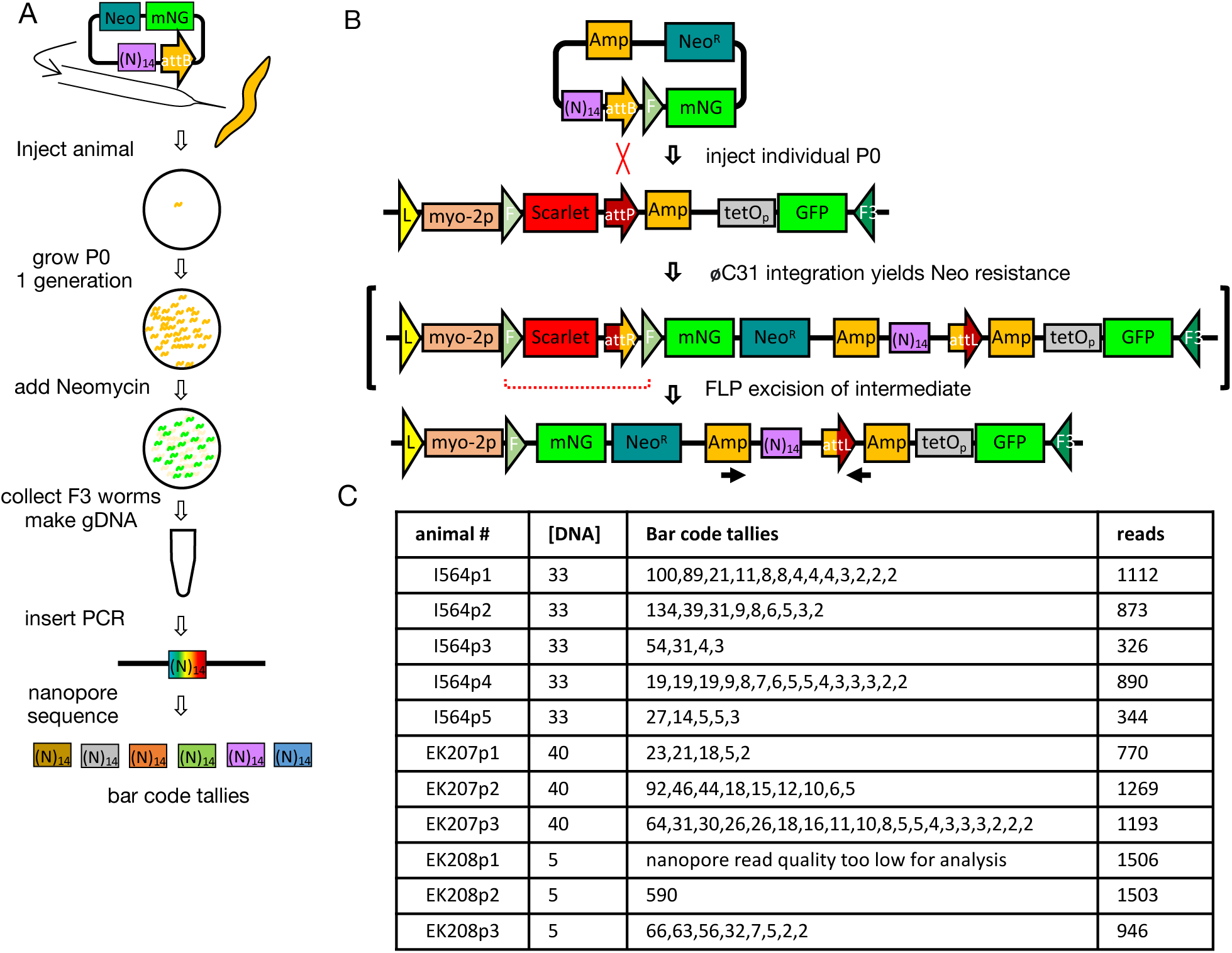
Quantification of RMI frequency using a bar code library. **A)** Schematic of the methodology used to quantify RMI efficiency. A bar-coded pool of ∼10,000 plasmids was created by amplifying an empty RMI integration plasmid using two oligos appended with 5’ (N)_7_, then re-circularizing the amplified product to create plasmids with (N)_14_ as the insertion. The pool of plasmids was injected into 11 individual animals at differing concentrations. On day 3 neomycin was added to select for insertions. Six days after injection the plates all contained many animals expressing nls-mNG in the pharynx. These plates were allowed to starve, then the animals on each plate were collected *en masse* and genomic DNA was prepared from each individual plate. PCR was performed across the bar code using an oligonucleotide pair specific for insertion events and the PCR product was sequenced using nanopore technology. **B**) Detailed schematic of the RMI integration performed showing the position of the oligonucleotides used to quantify integration frequency. *attB* and *attP* sites are shown as orange and red arrows and the recombined *attL* and *attR* sites as dual color arrows based on the recombination event. Yellow triangles labeled ‘L’ represent *loxP* sites, the cantaloupe triangle labeled ‘L2’ represent *lox2272* sites, and light and dark green triangles labeled ‘F’ and ‘F3’ represent *FRT* and *FRT3* sites, respectively. The orientation of the triangle represents the orientation of these recombination sites in the construct. **C**) Table of bar code tallies obtained from nanopore sequence analysis of the progeny of individual injected animals. The total number of reads does not equal to bar code tallies because of the high error rate of nanopore sequencing. Strain injected NM6456 *jsSi2284 II; jsSi1784 IV*. Plasmid injected NMp5004.

### Pseudo *att* sites are not a significant source of integration events in *C. elegans*

In most systems where phiC31-dependent integration has been performed, integration occurs at both introduced *att* sites and *pseudo*-*att* sites in the genome that have some similarity to consensus *att* sites (Thyagarajan et al., 2001; Combes et al., 2002; Groth et al., 2004; Allen and Weeks, 2005). To test if the *C. elegans* genome contains *pseudo-attP* or *pseudo-attB* sites I first constructed integration vectors containing a *FRT* site, a self-excisable *sqt-1(e1350) Hyg^R^* cassette, a MCS, and either an *attB* site or an *attP* site. After inserting a promoter FP transcription unit into the vectors, the integration plasmids were injected into animals expressing both FLP and phiC31 in the germline (**Fig. 8A**). A FLP expressing background was used to prevent array. In a first set of experiments, Rol animals present on day 3 were picked and screened on day 6 for transmission. In the case of the *attB* plasmid injections, no transmission was observed from 189 Rol animals derived from injection of 28 P0 animals. In the case of the *attP* plasmid injections, 1 insertion was obtained from 168 Rol animals derived from injection of 32 P0 animals (**Fig. 8B**). The single insertion was molecularly characterized and determined to be an integration at one of the *lox511* sites of the plasmid into a pseudo *lox511* site on Chr I (**Fig. 8C**). This insertion is presumed to have been mediated by leaky Cre expression from the *hsp-16.1* promoter present in the SEC of the plasmid. The first set of experiments was performed without Hyg^R^ selection to eliminate the possibility that integration events might occur in the germline of mosaic F1 animals that would not be Hyg^R^.

**Figure 8.**
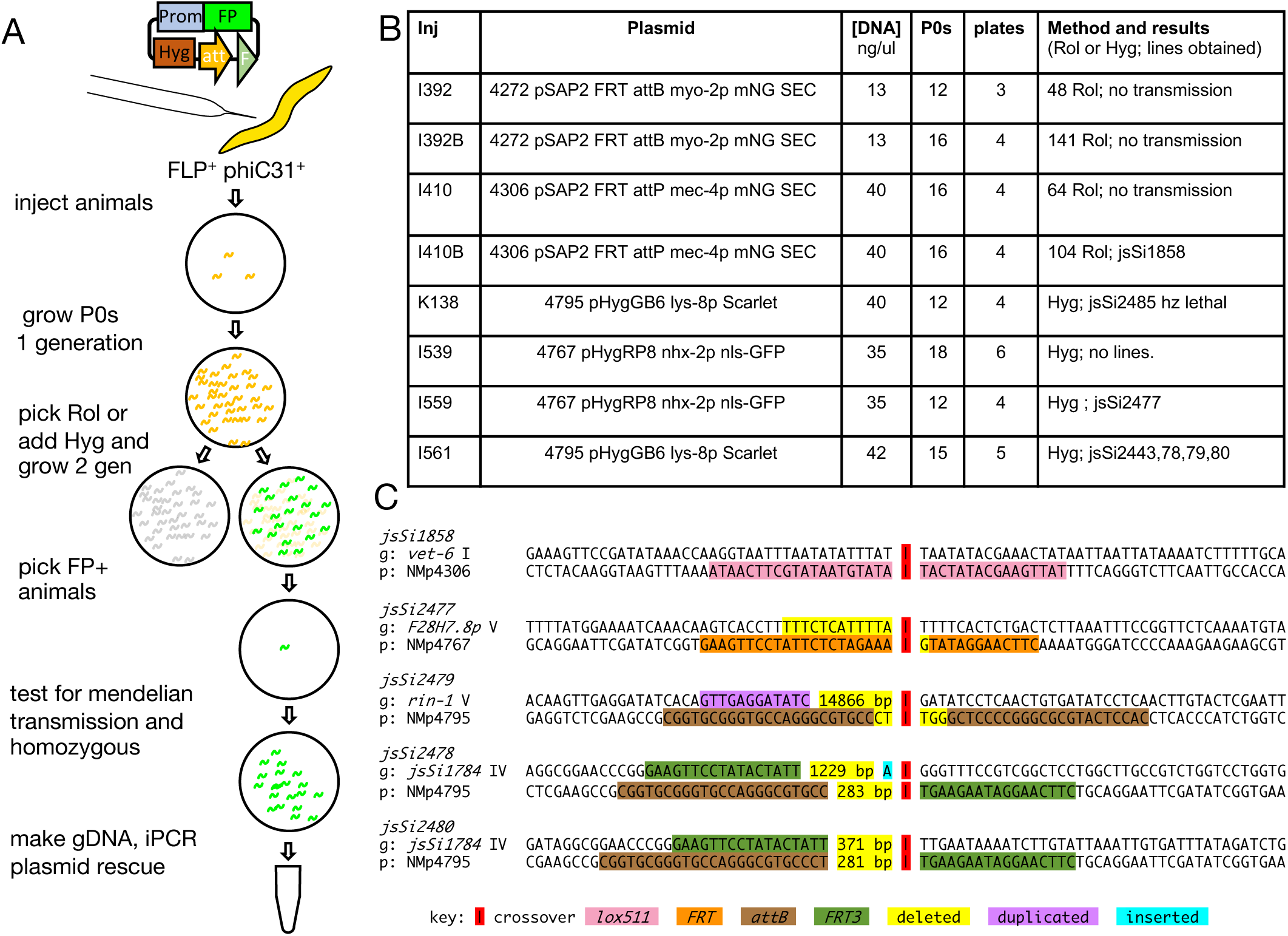
Test for the presence of pseudo-*att* sites in *C. elegans*. **A)** Schematic of the methodology used to determine the frequency of pseudo-*att* sites in *C. elegans*. In brief, young adult worms expressing both phiC31 and FLP but not containing an integrated *att* site were injected with a plasmid containing both an *att* site, visual and selection markers, and a *FRT* site. One generation after the injection, hygromycin was added and two generations later (F3 generation), individual hyg^R^ FP^+^ animals were cloned and characterized for Mendelian segregation of the fluorescent reporter. Genomic DNA was prepared from transmitting lines, then the insertion was characterized using a combination of inverse PCR and plasmid rescue. A similar strategy using a screen for Rol animals that transmit rather than a HygR selection was also performed. **B)** Table summarizing the results from eight injection sessions aimed at identifying *pseudo-att* sites. The first four injections were performed using a screen for transmission of a Rol phenotype. The second four injections were performed using Hyg^R^ to identify insertion events. **C)** Molecular characterization of the site of insertion of the injected plasmids. Shown is the sequence of the genomic site of each characterized integration (g) and the site of the integration in the plasmid (p). The site of the cross over is marked with a red bar (|). The sites of relevant recombinase sites as well as sequences inserted, deleted, or duplicated are shown in different colors as outlined in the key.

In a second set of experiments, integration plasmids containing a *Hyg^R^* gene, *loxP, FRT,* and *FRT3* sites, a promoter FP expression cassette, and either an *attB* site or an *attP* site, but no *Cre* gene, were injected into animals expressing both FLP and phiC31 (**Fig. 8A**). Hygromycin was added on day 3 and Hyg^R^ resistant animals were characterized. Among the progeny of 24 P0 animals injected with the *attP* plasmid, one integrated line was isolated (**Fig. 8B**). Molecular characterization revealed the insertion occurred by recombination between the *FRT* site and a pseudo-*FRT* site on Chr V (**Fig. 8C**). In the case of the *attB* plasmid, 33 P0 animals were injected and I obtained 5 insertions (**Fig. 8B**). One behaves genetically as an X-linked recessive lethal and a second could not be molecularly defined using either plasmid rescue or iPCR. The third integrated at the plasmid *attB* site while deleting 14.86 kb of the *rin-1* gene on Chr V (**Fig. 8C**). The last two integrated at the *FRT3* site of the *jsSi1784* source of phiC31, resulting in the deletion of sequences between the *attB* site and distinct “pseudo *attP”* sites adjacent to the transgenes. One of these two deleted 371 bp and the other 1.23 Kb. The latter deleted into *dpy-13* yielding a Dpy phenotype for the insertion. Both insertions were genetically mapped to confirm the position of the insertion. These data are consistent with these two insertions occurring by FLP mediated insertion at the FRT3 site followed by an *attB pseudo-attP* excision event (**Fig. S13**). In totality, these findings suggest that the frequency of pseudo *att* sites in the *C. elegans* genome is much lower than the rate of insertion into transgenic *attB* or *attP* sites. I conclude that integration into pseudo-*att* sites is unlikely to be a significant background in RMI experiments and also unlikely to disrupt RMHE recombination.

## Discussion

Bipartite expression systems are widely used in multiple model systems but have largely been ignored in the *C. elegans* community. Here I describe two new techniques to facilitate the creation and manipulation of bipartite reporters both of which take advantage of the robust activity of phiC31 recombinase. RMHE allows one to create novel *cis*-linked combinations of drivers and reporters by recombining the two components onto a single chromosome. RMI provides an alternative approach to creating *cis*-linked driver-reporter combinations by directly integrating into *att* tagged reporters or drivers. Although both techniques have been developed for manipulation of bipartite tools, RMHE and RMI techniques could also be used to catalyze similar chromosome exchange and integration events in other contexts. In particular, RMI-based transgenesis could relatively straightforwardly be developed into a general transgenesis technique that could be at least an order of magnitude more efficient than either CRISPR or RMCE based approaches.

### Strengths and limitation of RMHE

RMHE permits one to leverage the benefit of combinatorial power of bipartite reporter systems while simultaneously maintaining the simplicity of single locus genetics. However, there are costs associated with this approach. Specifically, for distinct driver and reporter transgenes to be RMHE compatible they must all be tagged in a similar manor and integrated at the same position. In developing the techniques, different orientations of driver and reporter combinations were tested which impose significant constraints on how drivers and reporters can be oriented with respect to each other (**Fig. 4G-J, S7 G-J**). Even after imposing these constraints, there are still many ways of creating tools that are poorly compatible, and only a few ways to make them optimally compatible. Novel tool development within a lab setting will likely coalesce around specific design. However, a community level commitment to tool development will likely be needed to extract the maximal power out of the approach. As a first step, I encourage the community to independently assess the utility of the approach.

### Strategies for identifying RMHE recombinants

Although RMHE is relatively efficient and often recombined chromosomes can easily be identified in the progeny of a single animal, but in some cases the identification of recombinant progeny can be challenging. Specifically, the recombinant chromosome needs to be distinguished from the *trans*-heterozygote. In developing the approach, I have settled upon several strategies that are efficient in identifying animals carrying the desired event. If the recombinant chromosome expresses a signal that can be visualized, then a reliable approach is to cross a male expressing an easily detected transgene that can be distinguished from the expected signal from the recombinant chromosome to animals heterozygous for the two unrecombined chromosomes (and expressing phiC31). Cross progeny can then be screened directly for the recombinant chromosome (**Fig. S8A**). In cases where rRMCE was used to create reporters and drivers expressing *cis*-linked markers, self-progeny carrying the recombinant chromosome can be distinguished from *trans*-heterozygotes by the loss of one of the *cis* markers making outcrossing unnecessary (**Fig. S8B**). The most challenging situation involves creating recombinant chromosomes that do not express a reporter that can be easily visualized. One such situation is the creation of cell specific GFP1-10 expressing lines for NATF. To simplify screening for such recombinants, a *tetO 7X tomm-20-GFP11* transgene *cis* marked with *myo-2p::nls-GFP* was created. Using males carrying this GFP11 transgene, recombinant GFP1-10 chromosomes can be identified as *myo-2 nls-GFP* positive cross progeny that express mitochondrial targeted GFP in the cell type of interest (**Fig. S8C**). Similar strategies can be implemented for other types of reporter constructs.

### Other uses for RMHE

Many transgenes have been inserted at a few well characterized loci; notably at mosSCI loci such as *ttTi5605* II and *cxTi10882* IV. RMHE offers a relatively simple way of utilizing these pre-existing well-vetted transgenes to create more complex ones. Introducing either an *attP* site 5’ of the insertion or an *attB* site 3’ of the insertion creates a RMHE compatible reagent that can be recombined with any new transgene created at the same site using rRMCE and the appropriate RMHE-compatible vector. Thus, in theory any two existing transgenes integrated at the same genomic position can be combined if they are modified using CRISPR methodology to contain the appropriate *att* site. However, it should be noted that recombining two transgenes can result in unexpected interactions between the insertions.

### Navigating the interactions in components of complex transgenes

The concept of creating strains with multiple linked reporters and effectors to simplify their use in both genetic analysis and exploratory genetic screens is appealing and potentially very powerful. However, at least in developing tools in my lab, it has become clear that interactions between promoters among closely spaced transcriptional units are common and unpredictable (Nonet, 2023; Knoebel et al., 2024). Furthermore, it is often difficult to nail down the cause of these effects. This can and will make development of these tools challenging. The identification of sequences that act as insulators could provide key resources for building such complex tools, though if such elements exist in *C. elegans* they will likely will act through distinct mechanisms than those previously defined in other systems (Heger et al., 2009). RMHE also offers a potential means of avoiding crosstalk issues by creating two integration loci separated by larger distances (e.g 10s to 100s of Kbs). Such transgenes will still be virtually completely linked (m.u.<.01 in most genetic context), but likely minimize crosstalk.

### Integration into the worm genome using RMI

phiC31 has previously been used to integrate plasmid-derived template DNA into the genome and specifically has been demonstrated to function in *C. elegans* to catalyze RMCE (Yang et al., 2022). In that approach phiC31 is expressed from a transgene flanked by *attP* sites and the RMCE event replaced the recombinase expressing cassette with sequences provided via injection of *attB* flanked targeting vector using *unc-119(+)* selection to identify recombination events. This strategy can potentially lead to insertion of multiple copies of the plasmid, though this did not appear to be common. Although a RMCE approach is not compatible with integration into RMHE-compatible driver and reporter lines, single site integration is compatible.

Adopting a RMHE strategy to creating bipartite driver-reporter transgenes yields a rich repertoire of driver and reporter lines that are tagged with an *att* site. I initially developed RMI to more rapidly develop novel reporters and drivers for this system. Development of novel lines often requires several rounds of tuning to optimize. Levels of expression are often inappropriate, and the functionality of a reporter can be influenced by the position of the marker and the size of the linker spanning domains. RMI provides a rapid approach to test new drivers and reporters by directly integrating into a complementary strain. This greatly increases the time efficiency of performing optimization. Furthermore, the resulting product is also a fully functional tool, though it does not produce an *attB* or *attP* tagged line to utilize to create other lines using RMHE. It remains possible that a phiC31 disintegrase mutant could be used to convert RMI constructs into new driver and reporter tools (Knapp et al., 2015).

### RMI is efficient

RMI appears to be surprisingly more efficient than RMCE. There are multiple possible reasons for this. First RMI only requires a single recombination event to occur while RMCE requires two. Second, phiC31 could simply be a more potent recombinase. phiC31 is capable of RMHE, but FLP is not (**Fig. 1**, **2**). One interpretation of this data is that phiC31 is a more efficient recombinase in the context of the *C. elegans* genome. This efficiency could be the result of the potency of the recombinase or the level of recombinase that is expressed in the worm germline. Regardless of the reason that RMI appears more efficient, the outstanding efficiency of RMI suggests that it may be worth further developing the system as a general highly efficient transformation technique.

### A diversity of potential RMI methods

A variety of approaches could be used to develop a general transgenesis pipeline taking advantage of RMI. The simplest approaches would only require introducing minimal *attB* or *attP* sites into the genome using an oligonucleotide templated CRISPR approach. These could include often used sites like ttTi5605 II and *cxTi10882* IV as well as sites at other desired locations. With these *att* sites in a FLP and phiC31 germline expression background, sequences can be introduced into a floxed HygR or NeoR vector, then introduced into the genome by injection (**Fig. S14A**). The selection cassette and vector could subsequently be excised by crossing through germline Cre lines (Nonet, 2023). One disadvantage of this approach is that the landing sites are not marked and thus more difficult to follow during crosses. Furthermore, the source of FLP and phiC31 must be provided in *trans* and removed after integration. Removing this source can be done in conjunction with removing the selectable marker. However, if RMI is being used as a rapid screen for the functionality of novel transgenes and this functionality can be assayed in the presence of the recombinase transgene and the selectable marker, then this approach will be very valuable.

Alternatively, more complex landing sites could be created that incorporate the recombinases analogous to how the these are incorporated into landing sites in the rRMCE approach (**Fig. S14B**). In this scenario, the sequence of interest would be integrated while simultaneously introducing a *cis*-linked marker to follow the insertion. The recombinase source would be excised during insertion, eliminating the requirement to outcross the novel transgene. The *cis* marker and/or selection cassette could still optionally be excised by outcrossing through germline Cre lines. Such an approach would provide much more flexibility in creating transgenes with distinct structure and simplify manipulation of landing sites by having the landing sites visually marked and encoding the recombinase. However, developing these landing site reagents would be significantly more complicated.

Another class of landing sites one might consider developing are landing sites that contain multiple *attB* or *attP* sites which would potentially provide the ability to perform reiterative insertions. Alternatively, a second *att* site could be co-inserted with the transgene providing an approach to create an insertion that contains a site to perform an additional integration. These approaches are most feasibility developed using an unlinked source of phiC31 and Cre. Hopefully the efficiency of phiC31 recombination will stimulate others to develop creative methods for creating complex transgenes.

### Stability of landing sites

While assessing the frequency of *pseudo-att* sites in the genome, two concerning insertion events, *jsSi2478* and *jsSi2480*, were identified. The most likely mechanism that led to these insertions is the excision of a small genomic region by a recombination between an integrated *attB* site and a *pseudo-attP* site. If such events occur at a reasonable frequency, then *attB* or *attP* sites introduced into the *C. elegans* genome and maintained in the presence of a germline source of phiC31 might not be completely stable. Similar concerns have been brought up in other systems that use phiC31 to create transgenes including both mammalian systems and Drosophila. Analysis of mammalian tissue culture lines with insertions into *pseudo-att* sites indicate that the vast majority are simple insertions, with only a minority (<10%) being associated with deletions or translocations (Chalberg et al., 2006). In these experiments transfection with a pCMV-int construct was used and phiC31 levels were likely transiently very high which could increase the frequency of unusual events. Mice that express phiC31 constitutively in the germline develop normal and are fertile (Belteki et al., 2003). Similarly, *attP* landing sites can be maintained in the presence of constitutive germline expression of phiC31 in Drosophila (Bischof et al., 2007). My lab has also maintained multiple *att* site containing RMHE drivers and reporters in the presence of germline phiC31 for dozens of generations without disrupting the *att* sites. The ability to maintain many species that express phiC31 in the germline constitutively argues that the spurious recombination by phiC31 is rare. The two unusual events I characterized appeared at the frequency of 1 event in ∼30 animals injected whose progeny were kept in the presence Hygromycin selection for two generations. By contrast, RMI events occur at a frequency of ∼5-10/ animal. Thus, it seems that recombination between *att* sites and *pseudo-att* sites are likely very rare events at least in *C. elegans*.

### Multicopy integration

RMI would appear to be an ideal approach to creating multicopy integrated arrays since phC31 is capable of relatively efficiently recombining an *attB* and *attP* site on distinct chromosome sized DNA as demonstrated by RMHE. However, attempts to derive multicopy insertions have largely been unsuccessful. Co-injection of a mixture of an *att* containing plasmid and other marker containing plasmids all lacking FRT sites (to allow for efficient array formation) into many dozen animals harboring an *attP* landing site and expressing phiC31 only resulted in a handful of resistant animals that were exceedingly unhealthy. This could be because high copy number of a *hygR* gene is toxic, or that the center of Chr II is not a genomic position compatible with large insertions. Various derivatives of this type of experiment including injecting linear DNAs and spiking the mixture with a small fraction of FRT containing HygR plasmids also failed to isolate multicopy insertions except for a few rare examples. Nevertheless, further attempts are probably warranted. It may be that array formation is too rare in injections of individual animals and that strains carrying arrays will need to be isolated and subsequently crossed into phiC31 *att* landing site strains to drive integration of multi-copy arrays into the genome by recombination.

In summary, I have developed and demonstrated the utility of two new techniques for the manipulation and creation of transgenic animals in *C. elegans*. Although both techniques were designed specifically to facilitate the development and manipulation of bipartite driver-reporter transgenes, they also show promise for the creation of other types of transgenic animals. In particular, the efficiency of RMI raises the potential of utilizing *C. elegans* transgenesis not only as a method to create tools, but also directly as a discovery tool as it is used in yeast, bacteria and tissue culture.

## Methods

### Nomenclature

*C. elegans* RMCE insertions into a landing site locus (e.g., *jsSi1726*) should technically be called *jsSi1726 jsSi#* according to *C. elegans* nomenclature rules (Tuli et al., 2018), but were referred to in the paper as *jsSi#*. A list of strains and a list of transgene insertions are provided in Table S1. These lists use the a <{region reverse oriented} nomenclature to indicate that a section of the insertion is in the reverse orientation.

### *C. elegans* strain maintenance

*C. elegans* was grown on NGM plates seeded with *E. coli* OP50 on 6 cm plates. Stocks strains were maintained at RT (∼ 22.5°C). (r)RMCE experiments were performed at 25°C.

### Plasmid microinjections

Injections were performed as described in (Nonet, 2023). Most animals were only injected in a single gonad. DNAs were injected at ∼ 50 µg/ml in 10 mM Tris pH 8.0, 0.1 mM EDTA.

### Isolation of RMCE and RMI insertions

Insertions were created using RMCE, rRMCE and RMI. The RMCE *sqt-1* Rol screening strategy was performed as outlined in Nonet, 2020. For rRMCE using drug selection (Nonet, 2023) or RMI, injected P0 animals were pooled 2-3 per plate in most cases. For RMI injections used to quantify insertion frequency using nanopore sequencing, a single injected animal was placed on each plate. For drug selection, the following were added directly to worm plates 3 days after injection: Hyg^R^ selection −100 ul of 20 mg/ml hygromycin B (GoldBio, St. Louis, MO), Neo^R^ selection - 500 ul of 25 mg/ml G418 disulfate (Sigma, St. Louis, MO). Eight days (6 days for RMI) after injection L4 animals homozygous for insertions could usually be identified on injection plates. The integration plasmid and parental strain used to construct each novel transgenic insertion and the strains containing these alleles and all other strains used in the study are listed in Table S1.

### Characterization of RMCE, rRMCE and RMI insertions

The structure of all novel RMCE landing sites and some RMCE insertions were analyzed using long range PCR of genomic DNA as previously described (Nonet, 2020). Genomic DNA was isolated as described in **supplemental methods**. Except when otherwise noted, oligonucleotides NMo6563/6564 were used to amplify Chr I insertions, NMo3880/3884 or NMo3887/3888 for Chr II insertions, NMo3889/3890 for chr IV insertions, and NMo7280/7281 for Chr V insertions. Some RMCE insertions into landing sites were sequenced, but most were only analyzed by restriction digestion of PCR products to confirm the insertion structure. *pseudo-att* insertions were characterized by inverse PCR as described in (Nonet, 2020) and/or by plasmid rescue (**supplemental methods**).

### Plasmid and vector constructions

The term vector, rather than plasmid, was used to distinguish the parental integration vectors from integration plasmids that contain specific sequences inserted in the MCS of the vectors. All PCR amplifications for plasmid and vector constructions were performed using Q5 polymerase (New England Biolabs, Ipswich, MA). Most PCR reactions were performed using the following conditions: 98°C for 0:30, followed by 30 cycles of 98°C for 0:10, 62°C for 0:30, 72°C for 1:00/kb). PCR products were digested with *DpnI* to remove template if amplified from a plasmid, then purified using a standard Monarch (New England Biolabs) column purification procedure. The invitrogen 1 Kb ladder Plus Ladder (cat # 10787018) was used as a marker in figures presenting PCR analysis. Restriction enzymes (except for *LguI*), T4 DNA ligase, and polynucleotide kinase were purchased from New England Biolabs. Golden Gate (GG) reactions (Engler et al., 2008) were performed as described in Nonet, 2020 except that in some cases LguI (Thermo Scientific™, Waltam, MA) was used in place of *SapI*. The *E. coli* strain DH5α was used for all transformations. Sanger sequencing was performed by GENEWIZ (South Plainfield, NJ) and nanopore sequencing by Plasmidsaurus (Eugene, OR). Oligonucleotides were obtained from MilliporeSigma (Burlington, MA) and synthetic DNA fragments were purchased from Twist Biosciences (South San Francisco, CA). A detailed description of all constructs is provided in **supplemental methods**. The sequence of all plasmids, vectors and synthetic fragments and oligonucleotides is provided in **Table S1**.

### Microscopy

Screening of worms for fluorescence during the RMCE protocol was performed using a Ziess Stemi SV11 dissecting microscope outfitted with a Kramer M2 bio epifluorescence module with a 1.6X lens and Kramer Scientific 10X (n.a. 0.45) LWD objective for high power observation illuminated using a Lumencor (Beaverton, OR) Sola light source. For imaging or quantification of fluorescence, worms were mounted on 2 % agarose pads in a 2 µl drop of 1 mM levamisole in phosphate buffered saline. 10 to 20 L4 animals were typically placed on a single slide. In cases where muscle was imaged, individual animals were placed in a drop of 8 ul of 1 mM levamisole in phosphate buffered saline and diluted 25 um diameter Polybead® Polystyrene beads (PolySciences Inc, Warrington, PA). After addition of a coverslip, the animals were rolled to a ventral or dorsal position by gently pushing the coverslip with a pipette tip. Animals were imaged using a 10X air (n.a. 0.45), 20X air (n.a 0.5) or 40x air (na 0.75) lens on an Olympus (Center Valley, PA) BX-60 microscope equipped with a Qimaging (Surrey, BC Canada) Retiga EXi monochrome CCD camera, a Lumencor AURA LED light source, Semrock (Rochester, NY) GFP-3035B and mCherry-A-000 filter sets, a Chroma (Bellows Falls, VT) 89402 multi-pass filter set and a Tofra (Palo Alto, CA) focus drive, run using Micro-Manager 2.0ß software (Edelstein et al., 2014). However, due to equipment failure, later images were taken after replacing the Retiga with a ToupTek MAX04BM sCMOS camera. The size of L4s often necessitated taking images with the animal in a diagonal position across the field. For figure panels such images were rotated leading to the presence of black corners in many panels. These black corners were adjusted to the mean intensity of a 3-pixel width line adjacent to the diagonal of each corner using a custom imageJ macro. Data plots were created using PlotsOfData (Postma and Goedhart, 2019).

## Data Availability

A full description of all oligonucleotides, plasmids, transgenes, and *C. elegans* strains created and used in this article are in Table S1. *C. elegans* strains to be sent to the Caenorhabditis Genetics Center are listed in Table S1. All other reagents are available upon request to Michael Nonet.

## Supplemental Materials

The manuscript contains 14 supplemental figures, two supplemental tables, a supplemental movie, and two supplemental text files named supplemental methods and supplemental figure legends.

## Acknowledgements

I thank Emma Knoebel for creating integration plasmids and isolating numerous transgenes, Dany Matar for creating several driver and reporter clones, Anna Brinck for comments on the manuscript, Jordan Ward, Matt Rich, and Han Wang and especially Tim Schedl for valued discussions about many aspects of developing these methods. Some strains were provided by the CGC, which is funded by NIH Office of Research Infrastructure Programs (P40 OD010440).

## Abbreviations

TRN: touch receptor neuron
CRISPR: Clustered Regularly Interspace Short Palindromic Sequences
RMCE: Recombination-Mediated Cassette Exchange
rRMCE: rapid RMCE
RMHE: Recombination-Mediated Homolog Exchange
RMI: Recombination-Mediated Integration
3’ UTR: three prime untranslated region
MT: microtubule
mNG: monomeric Neon Green FP
FP: Fluorescent Protein
GFP: Green FP
BFP: Blue FP
cyOFP: cyan-excitable Orange FP
SEC: Self-Excision Cassette

